# Workflows for rapid functional annotation of diverse arthropod genomes

**DOI:** 10.1101/2021.06.12.448177

**Authors:** Surya Saha, Amanda M Cooksey, Anna K Childers, Monica F Poelchau, Fiona M McCarthy

**Affiliations:** Boyce Thompson Institute, 533 Tower Rd., Ithaca, NY 14853 USA; School of Animal and Comparative Biomedical Sciences, 1117 E. Lowell Street, Tucson AZ 85721 USA; CyVerse, BioScience Research Laboratories, 1230 N. Cherry Ave., Tucson, AZ 85721 USA; USDA, Agricultural Research Service, Beltsville Agricultural Research Center, Bee Research Laboratory, 10300 Baltimore Avenue, Beltsville, MD 20705, USA; USDA, Agricultural Research Service, National Agricultural Library, 10301 Baltimore Avenue, Beltsville, MD 20705, USA

**Keywords:** Functional annotation, Gene Ontology, pathways, annotation, workflow, invertebrate

## Abstract

Genome sequencing of a diverse array of arthropod genomes is already underway and these genomes will be used to study human health, agriculture, biodiversity and ecology. These new genomes are intended to serve as community resources and provide the foundational information that is required to apply ‘omics technologies to a more diverse set of species. However, biologists require genome annotation to use these genomes and derive a better understanding of complex biological systems. Genome annotation incorporates two related but distinct processes: demarcating genes and other elements present in genome sequences (structural annotation); and associating function with genetic elements (functional annotation). While there are well established and freely available workflows for structural annotation of gene identification in newly assembled genomes, workflows for providing the functional annotation required to support functional genomics studies are less well understood. Genome-scale functional annotation is required for functional modeling (enrichment, networks, etc.) and a first-pass genome-wide functional annotation effort can rapidly identify under-represented gene sets for focused community annotation efforts. We present an open source, open access and containerized pipeline for genome-scale functional annotation of insect proteomes and apply it to a diverse range of arthropod species. We show that the performance of the predictions is consistent across a set of arthropod genomes with varying assembly and annotation quality.Complete instructions for running each component of the functional annotation pipeline on the command line, a high performance computing cluster and the CyVerse Discovery Environment can be found at the readthedocs site (https://agbase-docs.readthedocs.io/en/latest/agbase/workflow.html).

**Simple summary:** Genomic technologies are accumulating information about genes at a faster rate than ever before, and sequencing initiatives like the Earth Biogenome Project, i5k and Ag100Pest are expected to increase this rate of acquisition. However, if genomic sequencing is to be used for improvement of human health, agriculture and our understanding of biological systems, it is necessary to identify genes and understand how they contribute to biological outcomes. While there are several well-established workflows for assembling genomic sequences and identifying genes, understanding gene function is essential to create actionable knowledge. Moreover this functional annotation process must be easily accessible and provide information at a genomic scale to keep up with new sequence data. We report a well defined workflow for rapid functional annotation of whole proteomes to produce Gene Ontology and pathways information. We test this workflow on a diverse set of arthropod genomes and compare it to common arthropod reference genomes. The workflow we described is freely and publicly available via a web interface on CyVerse or as biocontainers that can be deployed scalably on local computing systems.

## Introduction

Over the past decade, rapid developments of sequencing technologies and assembly tools and algorithms have moved the bottleneck in genomics from data generation to inference of biological function. Model organism databases with sustained manual curation efforts have provided a source for homology [1,2] and - more recently - phylogeny-based [3] functional prediction for newly annotated gene sets. As we expand the sequencing efforts to organisms in hitherto poorly sampled branches of the eukaryotic tree of life [4], there is an increase in the number of novel proteins of unknown function and even identifying genes closely related to previously studied genes in other species can be problematic. While workflows have been developed to support genome assembly and gene identification, the process for understanding the function of resulting gene products is not as well documented.

Annotation spans two related but distinct processes in genomics: demarcating genes and other elements present in genome sequences (structural annotation); and associating function with genetic elements (functional annotation). Here, we focus on functional annotation of gene sets based on Gene Ontology (GO) terms and metabolic pathways. Genome-scale functional annotation is required for functional modeling (enrichment, networks, etc.) and a first-pass genome-wide functional annotation effort can rapidly identify under-represented gene sets for focused community annotation efforts.

High throughput functional annotation relies on transferring functional information to unannotated proteins based upon analysis of functional domains and sequence homology [5,6]. While different software packages have been applied to this process, the general approach to first-pass functional annotation is similar (Figure 1). Protein sets are scanned for motifs and domains using resources like Pfam [7] and InterPro [8,9] and mapped to Gene Ontology terms using GO supplied mapping files. In addition to identifying shorter motifs and domains, BLAST analysis of full length sequences can identify similar sequences which already have GO or pathway annotations linked to them. Examples of tools that rely on sequence similarity include GOanna [5], BLASTKoala [10] and Blast2GO [11]. More recently the GO Consortium started using phylogenetic relationships to transfer GO terms [3]. The advantage of this approach is that evolutionary relationships provide more reliable evidence for conserved function than sequence similarity; however this approach still relies on manual curation, which cannot keep pace with gene discovery from large scale genome sequencing projects. Each of these sequence-based approaches relies on transferring GO terms associated with a gene product in one species to a gene product in another species, and the best practice for transferring GO terms is to limit this process to GO terms assigned based upon direct evidence [12].

**Figure 1:**
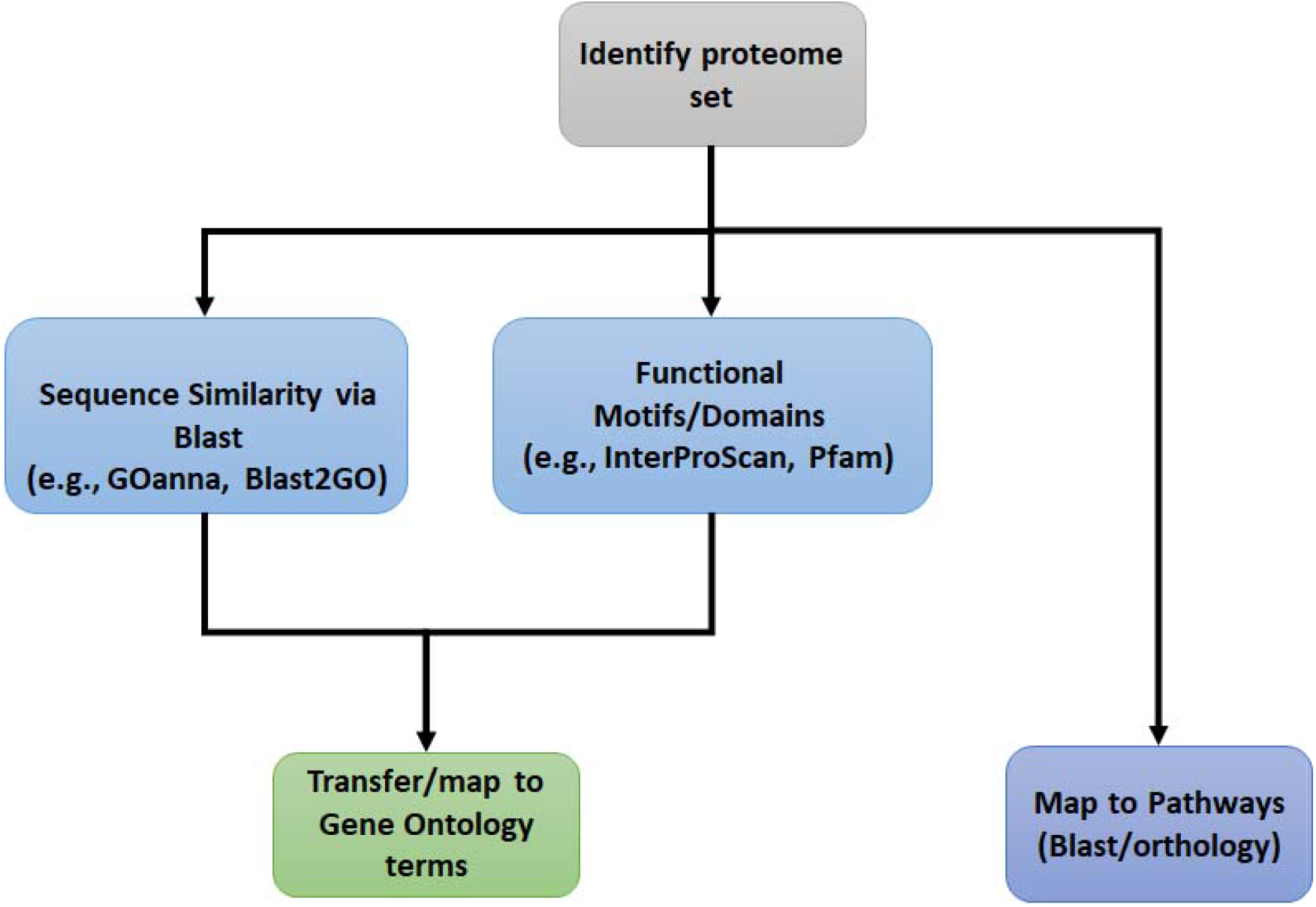
Generalized functional annotation workflow. The general approach for functional annotation is to combine GO annotations transferred on the basis of sequence homology (e.g., BLAST) with information about functional motifs (e.g., derived from resources such as PFAM). Gene products are mapped to metabolic and signalling pathways based upon sequence homology or orthology.

### Motivation

Many high-quality arthropod genomes are being generated, in particular by large-scale genome projects such as the Ag100Pest Initiative (http://i5k.github.io/ag100pest) and others under the Earth BioGenome Project umbrella [13]. These new genomes serve as community resources and provide the foundational information required to apply ‘omics technologies to a more diverse set of species. Genome assemblies need structural and functional annotations to ensure that these ‘omics approaches can be rapidly translated into biological information that provides a better understanding of the system being studied. The Gene Ontology Consortium [14], UniProtKB [15] and KEGG [15,16] resources generate and maintain functional annotations of many proteomes available in the sequence databases such as RefSeq and INSDC, and functional annotations produced by these initiatives are widely used and referenced by the scientific community. However, the process of manual curation of published papers is laborious and time consuming for model species where most publications are focused on gene function [17]. A rapid, first-pass functional annotation workflow quickly provides functional information to support genomic analyses and experimentation and ensures that ‘omics approaches can be interpreted to better understand a diverse range of biological systems.

AgBase [18] and the i5k Workspace@NAL [19] databases serve the arthropod genomics community by providing access and curation tools for arthropod proteomes and genomes, respectively. Here, we report the creation of containerized workflows to fill the need for high-throughput functional annotation of proteins from eukaryotic genome sequencing programs for the scientific communities that we support, as well as the arthropod genomics community at large. We test these workflows using twelve sequenced invertebrate genomes selected to span a broad range of invertebrate classes and to represent genomes with varying assembly quality (Table 3) and sequencing technologies used. These sequenced genomes are compared with three well studied invertebrate genomes, *Drosophila melanogaster, Apis mellifera (honeybee)* and *Tribolium castaneum* (red flour beetle). These workflows are also available on CyVerse to facilitate re-use [20,21] via a user-friendly web-based interface.

## Materials and Methods

### Functional annotation pipeline

Complete instructions for running each component of the functional annotation pipeline on the command line, a high performance computing cluster, or the CyVerse Discovery Environment can be found at the readthedocs site (https://agbase-docs.readthedocs.io/en/latest/agbase/workflow.html).

### Sequence similarity via BLAST: GOanna

GOanna [5] assigns GO terms based on sequence homology to specialized BLAST databases. These databases consist of proteins associated with GO terms, and grouped by phyla or taxonomic divisions (Table 1). The established best practice for transferring GO terms between similar sequences is to only transfer GO terms based upon experimental evidence codes, otherwise the risk of translative error increases substantially and functions inappropriate to the specie’s physiology are more likely to occur. GO uses several types of evidence to associate a GO term with a gene product: direct experimental evidence, phylogenetic relatedness and computational analysis. Transferring GO annotations based on experimental evidence codes is recommended to avoid inferring function based upon another inference. GOanna accepts a protein FASTA file as input and allows the users to set standard BLAST parameters (Supplementary Table 1). Since GOanna outputs results as a gene association file (GAF) file, it also requires users to provide information about the sequence source and species. Other information such as protein name is parsed from the FASTA header, and to ensure that it is correctly parsed from FASTA files generated by NCBI, an option to parse delimited sequence identifiers is also provided.

**Table 1.** GOanna version 2.2 databases. Databases are prepared from proteins that have GO annotations based upon taxonomic divisions. Protein numbers reported as of January 2019.

| Database Name | No. UniProtKB Proteins | No. in GOanna Db |
| --- | --- | --- |
| arthropod | 3,956,843 | 12,081 |
| bacteria | 28,660,834 | 12,748 |
| bird | 777,091 | 1,379 |
| fish | 1,505,807 | 12,478 |
| fungi | 7,614,812 | 13,718 |
| human | 161,566 | 21,125 |
| insecta | 2,883,005 | 11,886 |
| invertebrates | 8,409,505 | 20,741 |
| mammals | 1,836,549 | 42,966 |
| nematode | 1,541,602 | 4,941 |
| plants | 6,300,920 | 16,058 |
| UniProt-SwissProt | 50,258 | 72,337 |
| UniProt--TrEMBL | 4,720,107 | 57,834 |

### Functional motif analysis: InterProScan

InterPro ([8,9] is a database which integrates predictive information about protein function from a number of partner resources in the InterPro consortium. InterProScan ([8,9] is a software tool that accepts a FASTA file, identifies motifs and domains from InterPro protein databases (Table 2) and maps them to GO terms and pathways with a number of customizable parameters (Supplementary Table 2). Our dockerized implementation also performs checks to trim any unknown amino acids at the end of sequences including X’s as the inclusion of these often causes the platform to fail. It also removes the “*” symbol added by some translation software to denote a stop codon before running submitted protein sequences in parallel. Parallelization is an important consideration for scalability and utilization of high-performance computing resources. For those users with nucleotide sequences, documentation is provided for using TransDecoder [22] to translate open reading frames from transcripts. Moreover, many other options for translating sequences into proteins are also publicly available. The XML output from InterProScan is parsed to produce the output GAF file and report pathway information.

**Table 2:**
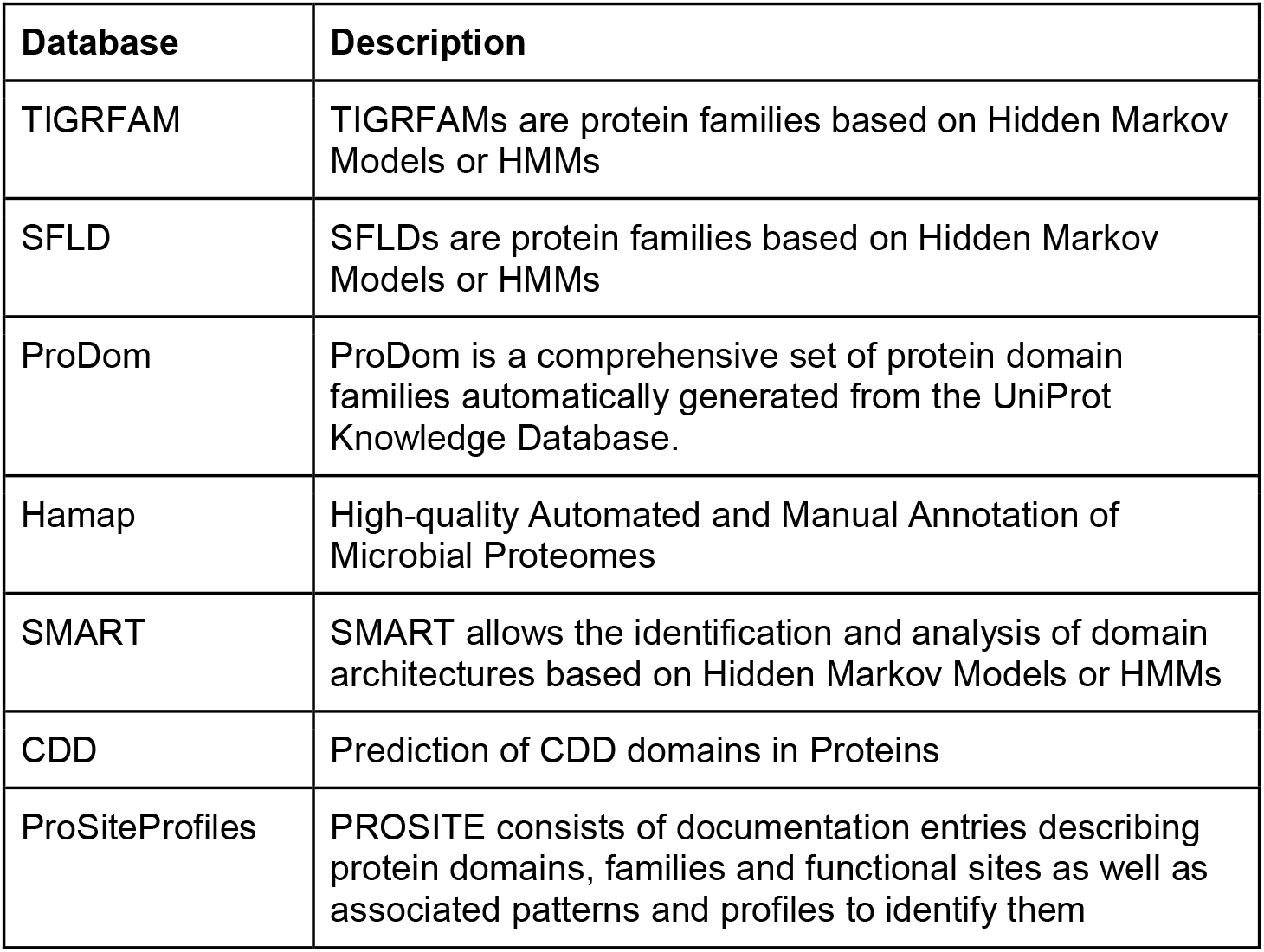
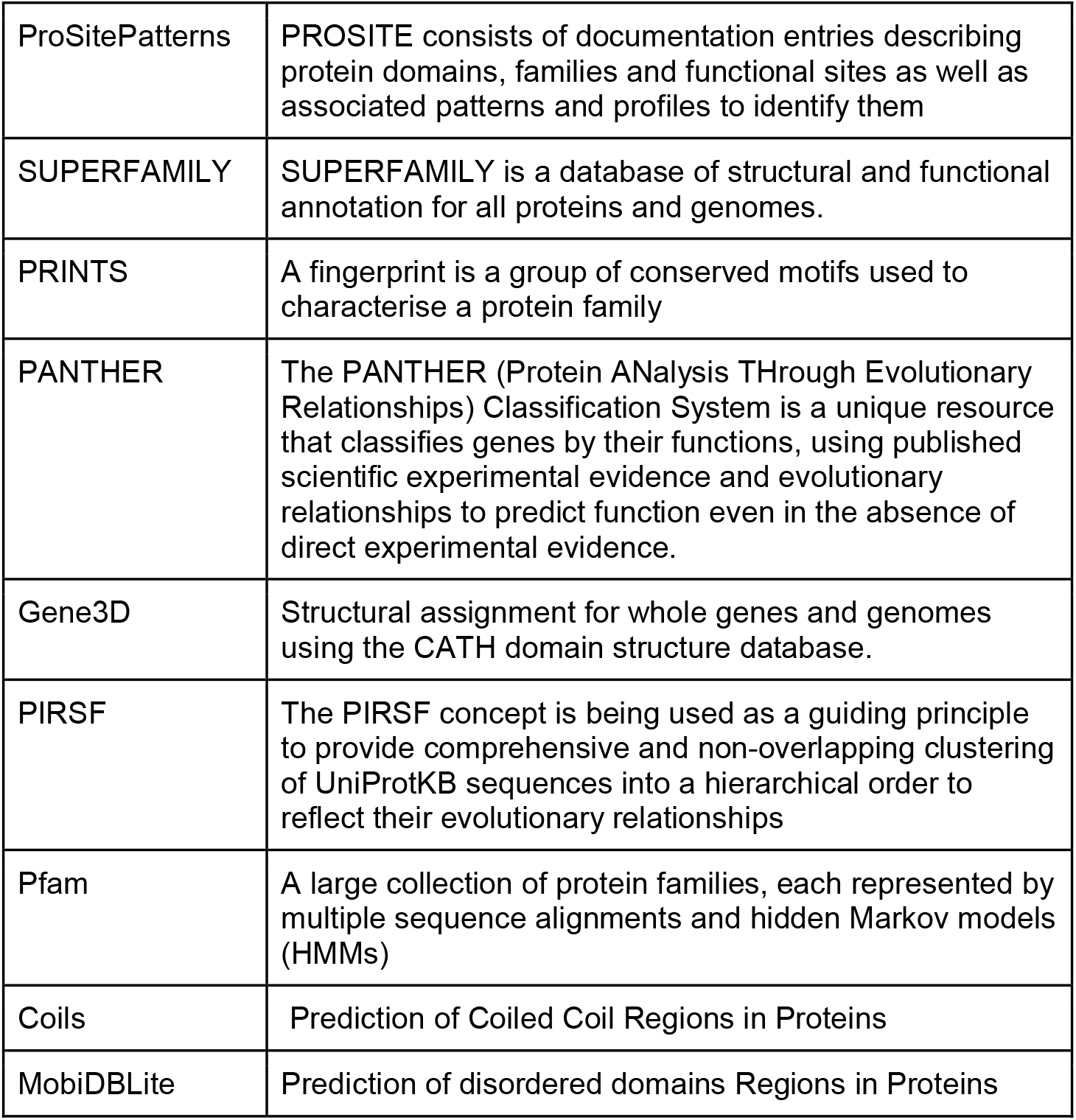
Databases used by InterProScan version 5.45-80 for annotation.

### Combining and QC of GO annotations

The GOanna and InterProscan containers both output a GAF, the standard file format for GO annotation data. This is a tab-separated file that can be easily combined, but for users who have large files that they cannot easily manipulate, the Combine GAFs tool we developed accepts multiple GAF files and combines them. Note that some users prefer to remove identical GO terms associated with the same protein by different software; since these GO terms are assigned by different methods and have different evidence codes, we do not remove these at this step.

In addition to combining GAF files, the GO annotation data can be assessed using the GO Annotation Quality (GAQ) Score [17]. GAQ is a quantitative measure of the quality of GO annotation of a set of proteins. GAQ scores include the breadth of GO annotation, the level of detail of annotation and the type of evidence used to infer the annotation. The scores generated can also be used to track changes in GO annotations over time. The GAQ tool determines the depth of each GO term and the rank of each evidence code associated with the annotation and returns a GAQ score as a product of depth and evidence code rank. The total GAQ score of each annotated gene product is calculated and a summary is generated showing the overall total GAQ scores, the number of gene products annotated and the average (mean) GAQ score of the whole protein set. We use the GAQ score to determine the value added to functional information, particularly when compared with well annotated model species such as Drosophila and to a lesser extent, *A. mellifera* and *T. castaneum*.

### Map to pathways: KOBAS

KEGG Orthology Based Annotation System (KOBAS) [23] assigns input proteins to known pathways in KEGG. It also includes a gene set enrichment function (Supplementary Table 3) to find statistically enriched genes in a disease or experimental condition with respect to the background of all annotated proteins in the organism. The pipeline consists of two modules:

- *Annotate:* This step assigns appropriate KEGG Ortholog (KO) terms for queried sequences based on a similarity search. It also assigns proteins to pathways from KEGG, Reactome and BioCyc.
- *Identify:* This performs an enrichment analysis compared to a background of the species’ gene set among the annotation results based on the frequency or statistical significance of pathways.

For annotating the gene products from a species, we use the *Annotate* module.

## Research Design and Method: Comparing Functional Annotation across Multiple Species

To test the usefulness of the functional annotation workflows, we selected a set of arthropod genomes (Table 3) with varying assembly quality and state of manual curation. This data set included several well studied arthropod genomes such as *Drosophila melanogaster, Apis Mellifera* and *Tribolium castaneum* for comparison. BUSCO [23,24] version 5.1.2 was used with the protein option and arthropoda_odb10.2019-11-20 database with 1013 markers to analyse all protein sets for completeness (Table 4).

**Table 3.** Arthropod genomes selected for this study and their assembly and annotation statistics. The test species are sorted by the scaffold N50 value.

| Species | Genome assembly accession | Genome assembly name | Contig N50 | Scaffold N50 | Annotation Name | Proteins | Proteins assigned GO terms | DOI |
| --- | --- | --- | --- | --- | --- | --- | --- | --- |
| *Apis Mellifera* (honey bee) | GCA_000002195.1 | Amel_4.5 | 5,832,476 | 13,619,445 | OGSv3.3 | 15,314 | 39.91% | NA |
| *Drosophila melanogaster* (fruit fly) | GCA_000001215.4 | DMEL_r6.36 | 21,485,538 | 25,286,936 |  | 30,724 | 59.42% | NA |
| *Tribolium castaneum* (red flour beetle) | GCA_000002335.3 | TCAS_5.2 | 73,049 | 4,456,720 |  | 18,534 | 44.98% | NA |
| *Latrodectus hesperus* (Western black widow spider) | GCA_000697925.1 | Lhes_1.0 | 2,223 | 13,889 | LHES-BCM_version_0.5.3 | 17,364 | 31.17% | 10.15482/USDA.ADC/1503795 |
| *Limnephilus lunatus* (caddisfly) | GCA_000648945.1 | Llun_1.0 | 2,103 | 54,650 | LLUN-BCM_version_0.5.3 | 13,292 | 55.76% | 10.15482/USDA.ADC/1503798 |
| *Oncopeltus fasciatus* (Large milkweed bug) | GCA_000696205.1 | Ofas_1.0 | 4,047 | 339,960 | oncfas_OGSv1.2 | 19,793 | 34.31% | 10.15482/USDA.ADC/1518752 |
| *Homalodisca vitripennis* (Glassy-winged sharpshooter) | GCA_000696855.1 | Hvit_1.0 | 4,857 | 512,049 | HVIT-BCM_version_0.5.3 | 33,019 | 38.00% | 10.15482/USDA.ADC/1410182 |
| *Eurytemora affinis* (calanoid copepod) | GCA_000591075.1 | Eaff_1.0 | 5,738 | 862,645 | EAFF-BCM_version_0.5.3 | 29,783 | 30.02% | NA |
| *Agrilus planipennis* (emerald ash borer) | GCA_000699045.1 | Apla_1.0 | 6,314 | 910,924 | APLA-BCM_v<br>ersion_0<br>.5.3 | 15,497 | 51.07% | 10.15482/USDA.A<br>DC/1503805 |
| *Copidosoma floridanum* (parasitoid wasp) | GCA_000648655.1 | Cflo_1.0 | 14,521 | 1,037,125 | CFLO-BCM_v<br>ersion_0<br>.5.3 | 19,869 | 34.14% | 10.15482/USDA.A<br>DC/1503793 |
| *Athalia rosae* (turnip sawfly) | GCA_000344095.1 | Aros_1.0 | 51,418 | 1,366,867 | AROS-BCM_v<br>ersion_0<br>.5.3 | 22,213 | 57.05% | 10.15482/USDA.A<br>DC/1459565 |
| *Ceratitis capitata* (Mediterranean fruit fly) | GCA_000347755.2 | Ccap_1.1 | 45,879 | 4,118,346 | Ccap-OGSv1 | 12,318 | 55.75% | NA |
| *Cimex lectularius* (Cimicidae bed bug) | GCA_000648675.1 | Clec_1.0 | 23,511 | 7,172,596 | Clec-OGSv1.2 | 14,212 | 49.42% | NA |
| *Varroa destructor* (parasitic mite) | GCA_002443255.1 | Vdes_3.0 | 201,886 | 58,536,683 | NCBI<br>Varroa<br>destruc<br>tor<br>Annotat<br>ion<br>Releas<br>e 100 | 30,221 | 53.60% | NA |
| *Diaphorina citri* (Asian citrus psyllid) | NA | Version 3 | 749,525 | 40,596,296 | OGSv3 | 19,049 | 59.30% | 10.1101/869685 |

**Table 4.** Arthropod genomes selected for this study and their BUSCO completeness statistics. The test species are sorted by the BUSCO completeness score. BUSCO version 5.1.2 was used with the protein option and arthropoda_odb10.2019-11-20 database with 1013 markers.

| **Species** | **Complete** | **Complete single-copy** | **Complete Duplicated** | **Fragmented** | **Missing** |
| --- | --- | --- | --- | --- | --- |
| *Drosophila melanogaster* (fruit fly) | 99.90 | 53.3 | 46.6 | 0 | 0.1 |
| *Athalia rosae* (turnip sawfly) | 99.70 | 68.9 | 30.8 | 0 | 0.3 |
| *Ceratitis capitata* (Mediterranean fruit fly) | 98.40 | 97.5 | 0.9 | 0.4 | 1.2 |
| *Tribolium castaneum* (red flour beetle) | 98.40 | 93.1 | 5.3 | 1.2 | 0.4 |
| *Apis Mellifera* (honey bee) | 97.40 | 96.9 | 0.5 | 1.5 | 1.1 |
| *Varroa destructor* (parasitic mite) | 95.90 | 43.1 | 52.8 | 0.7 | 3.4 |
| *Cimex lectularius* (Cimicidae bed bug) | 95.30 | 93.5 | 1.8 | 2.5 | 2.2 |
| *Copidosoma floridanum* (parasitoid wasp) | 93.70 | 92.5 | 1.2 | 2.9 | 3.4 |
| *Agrilus planipennis* (emerald ash borer) | 90.90 | 89.1 | 1.8 | 4.6 | 4.5 |
| *Diaphorina citri* (Asian citrus psyllid) | 87.10 | 55.9 | 31.2 | 2.8 | 10.1 |
| *Oncopeltus fasciatus* (Large milkweed bug) | 72.90 | 70.8 | 2.1 | 21.4 | 5.7 |
| *Eurytemora affinis* (calanoid copepod) | 57.50 | 55.9 | 1.6 | 20 | 22.5 |
| *Homalodisca vitripennis* (Glassy-winged sharpshooter) | 55.90 | 54.2 | 1.7 | 32.5 | 11.6 |
| *Limnephilus lunatus* (caddisfly) | 42.40 | 41.4 | 1 | 28.1 | 29.5 |
| *Latrodectus hesperus* (Western black widow spider) | 31.40 | 30.6 | 0.8 | 26.9 | 41.7 |

Proteome sets for each of the species were downloaded from NCBI and functionally annotated using the workflow described above. For GOanna, we used the invertebrate reference databases and only the GO terms with experimental evidence were assigned (-b). Custom BLAST parameters included a BLAST identity (-g) and query coverage (-q) cutoff of 70% with a maximum number of gap opening size (-k) of 9 to account for insertion or deletion of short peptides. Ideally the query and BLAST match should be of identical length, but we allowed for some flexibility (-r 1.2) to account for natural diversity and potential assembly or annotation errors.
InterProScan was run to identify InterPro domains, GO terms and pathways for the input proteins (-g -l -p -c) and we used all the databases in order to extract the maximum amount of information possible. A single, comprehensive GAF was obtained by combining the results from GOanna and InterProScan. The same protein sets were then run through KOBAS [23] to annotate pathways. The KOBAS Annotate tool (-a) used the Drosophila reference proteins (-s dme). The input data type has to be specified (-t fasta:pro).

## Results and Discussion

### Installation & Runtime considerations

The memory usage and runtime of the containers described here scales with the size of the protein set with the exception of InterProScan. The large number of databases (Table 2) that have to be searched for matches for each protein sequence increases the runtime and memory usage for even small data sets. The scalability of InterProScan has been improved with data and compute parallelization. The input proteins are split into sets of 1000 sequences for parallel processing, but the time required for loading and searching all the 16 databases is still significant. Another factor to consider is the increasing size of databases; new updates will only increase these requirements in the future. Therefore, we recommend that the InterProScan container be run on a high-performance computer like a cluster or a server with at least 256Gb of RAM and 500Gb of disk space. The documentation for this workflow (https://agbase-docs.readthedocs.io/en/latest/agbase/workflow.html) includes instructions on executing the containers with Singularity if Docker containers are not permitted due to security restrictions. The GOanna and KOBAS containers can be set up on desktop grade computers.

### Parameter optimization

Like all workflows, parameter optimization is a key part of ensuring quality results. Here we discuss the parameter optimization process for this workflow across a diverse range of arthropod genomes for new users to consider when applying this workflow to their own data sets. For the GOanna tool, the key optimization parameters are the selection of the database and the standard BLAST parameters. Many users prefer to do an initial BLAST search against a comprehensive database (e.g., NCBI nr or UniProt-SwissProt databases) to identify the most similar known sequence. While we include the UniProt SwissProt and TrEMBL database as options for GOanna, we note that the databases GOanna uses are not meant to be comprehensive but rather a subset of proteins that have been assigned GO terms. Moreover, given that searching against larger databases increases the probability of finding spurious matches, we recommend using the phyla specific database most relevant for your dataset and supplementing the output of GOanna matches with InterProScan results. To ensure high quality results, BLAST parameters should be optimized. While many analyses report optimizing BLAST solely on the E-value, this varies based upon database size. To determine BLAST parameters we randomly selected three sets of 1,000 sequences from each of the proteomes and manually reviewed the results of alignments from BLAST run with default parameters. The most common error when these sets were re-run with more stringent E-values was the identification of short, perfect matches (E-value = 0) that had low query coverage (e.g., less than 50%). To consistently return good matches from a broad range of protein sequences from all the proteomes used in this study, we used cut-offs of 70% identity and 70% coverage for the BLAST parameters.

Unlike GOanna which is BLAST-based, InterProScan searches for near perfect matches to short motifs and domains [9]. A key consideration for running InterProScan is to decide which databases should be searched. Some users prefer to analyze CDD or PFAM directly and both of these databases are included in the InterPro analysis. Since the computing requirements of InterProScan are considerable, these requirements could be reduced by searching fewer databases. While our workflow is deliberately designed to accept proteins, InterProScan can accept nucleotide sequences and translate them prior to searching the protein databases. Our initial tests indicated that submitting nucleotide sequences to InterProScan resulted in many more motif matches, but similar GO annotations (results not shown). Closer inspection revealed that the translation step produced large numbers of peptides but many did not match the known peptides produced from the mRNA sequence used as input. Therefore we recommend a separate translation step and submitting protein sequences to InterProScan.

To rapidly provide pathway annotations for arthropod gene products, we utilized the KEGG system which maps genes to pathways based upon sequence homology, creating KEGG Ortholog (KO) sets for different species. Since the KOBAS annotate tool takes a sequence file and uses BLAST to associate KEGG pathways with these sequences, parameter optimization requires the selection of the database to search against (e.g., “KO” for all orthologous proteins or “dme” to restrict to only Drosophila proteins) as well as standard BLAST parameters. The parameters (-e -r -C -z, designated by * in Table 6) denoting E-value, rank, subject coverage and orthologs for cross-species annotation can be modified to increase stringency when transferring annotation from the selected model species (-s). We note that the BLAST parameters required for this process may differ from GOanna because the two BLAST-related tools use different search databases.

### Overall summary of functional annotation of selected genomes

#### Genome assembly

To test our functional annotation workflow, we selected twelve arthropod genomes, four of which were community curated. The genomes were selected to represent a range of assembly quality and a diverse set of arthropod species. These twelve genomes were supplemented with three well-studied arthropods (a reference set): *Drosophila melanogaster* (fruit fly), *Apis mellifera* (honey bee) and *Tribolium castaneum* (red flour beetle) from the Orders Diptera, Hymenoptera and Coleoptera, respectively. We note that all of these species have been assembled, annotated, and the proteomes are considered mostly complete with BUSCO completeness scores ranging from 31 to 99% (Table 4). The genome assemblies for the selected species varied in contiguity and quality with scaffold N50s ranging from 13.8 kb to 58.5 Mb (Table 3). Another metric of interest for quantifying the quality of the assembly before scaffolding is contig N50 that ranged from as low as 2.2 kb for genomes assembled with Illumina paired-end and mate-pair reads to 749.5 kb for genomes assembled with PacBio long-read technology (Table 3). Please note that assemblies with low contig N50 but comparatively high scaffold N50 can have large gaps filled with unknown (N) nucleotides.

The proteome sets we used ranged from 12,318 - 33,019 proteins (Table 3). We examined the proportion of these proteins that were annotated with GO data, and were also interested in determining what BLAST-based analyses contributed to this GO annotation compared to the motif-based InterProScan annotation. Overall, GO annotation ranged from 30-60% of the protein set, with an average of 45% including the reference genomes. Notably, other species were able to achieve the same rates of GO annotation as the reference gene sets, indicating that the workflow performs as expected. We also wanted to evaluate if assembly contiguity (contig and scaffold N50) and gene space completeness corresponded to coverage of functional annotation for the proteome. This was not always the case as 44.6% of the proteins from *L. lunatus* (caddisfly) were associated with GO terms but the assembly only has a scaffold N50 of 54.6 kb and a contig N50 of 2.1 kb. The gene space for caddisfly is relatively incomplete at 42.4 with low duplication (Supplementary Figure 1 and Table 4). On the other end of the spectrum, the hymenopteran *C. floridanum* (parasitoid wasp) has a contig and scaffold N50 of 14.5 kb and 1 Mb, respectively, but only 34.1% of its proteins have GO terms associated with them. The other hymenopteran in the test set, *A. rosae* (turnip sawfly) has a better GO term coverage of 57.05%, but it also has a more contiguous genome with a contig and scaffold N50 of 51.4 kb and 1 Mb, respectively. Both *A. rosae* (turnip sawfly) and *C. floridanum* (parasitoid wasp) have comparable BUSCO completeness metrics (99.7% and 93.7%), but duplication in the gene space is higher at 30.8% in *A. rosae* compared to only 1.2% in *C. floridanum*.

#### Gene Ontology Annotation

BLAST-based GO annotation assigned markedly fewer GO terms (accounting for at most only 4.09% of assigned annotations in caddisfly) (Table 5). However, the value of the GO annotations added by BLAST-based tools like GOanna is disproportional to the quantity of GO added by these tools. We measured the value of the GO terms assigned to gene products using the GO Annotation Quality (GAQ) Score [9,17]. The average GAQ score for GO terms assigned by BLAST using GOanna was 142.02 while the average GAQ score of GO terms assigned by InterProScan based on motif search was 34.84. The GAQ score of the Drosophila functional annotation downloaded from European Bioinformatics Institute (EBI) [25], which included manual annotation, had a much higher GAQ score of 243.68 as it included evidence codes for manual functional annotation which are weighted higher than sequence similarity based GO term assignment.

**Table 5.**
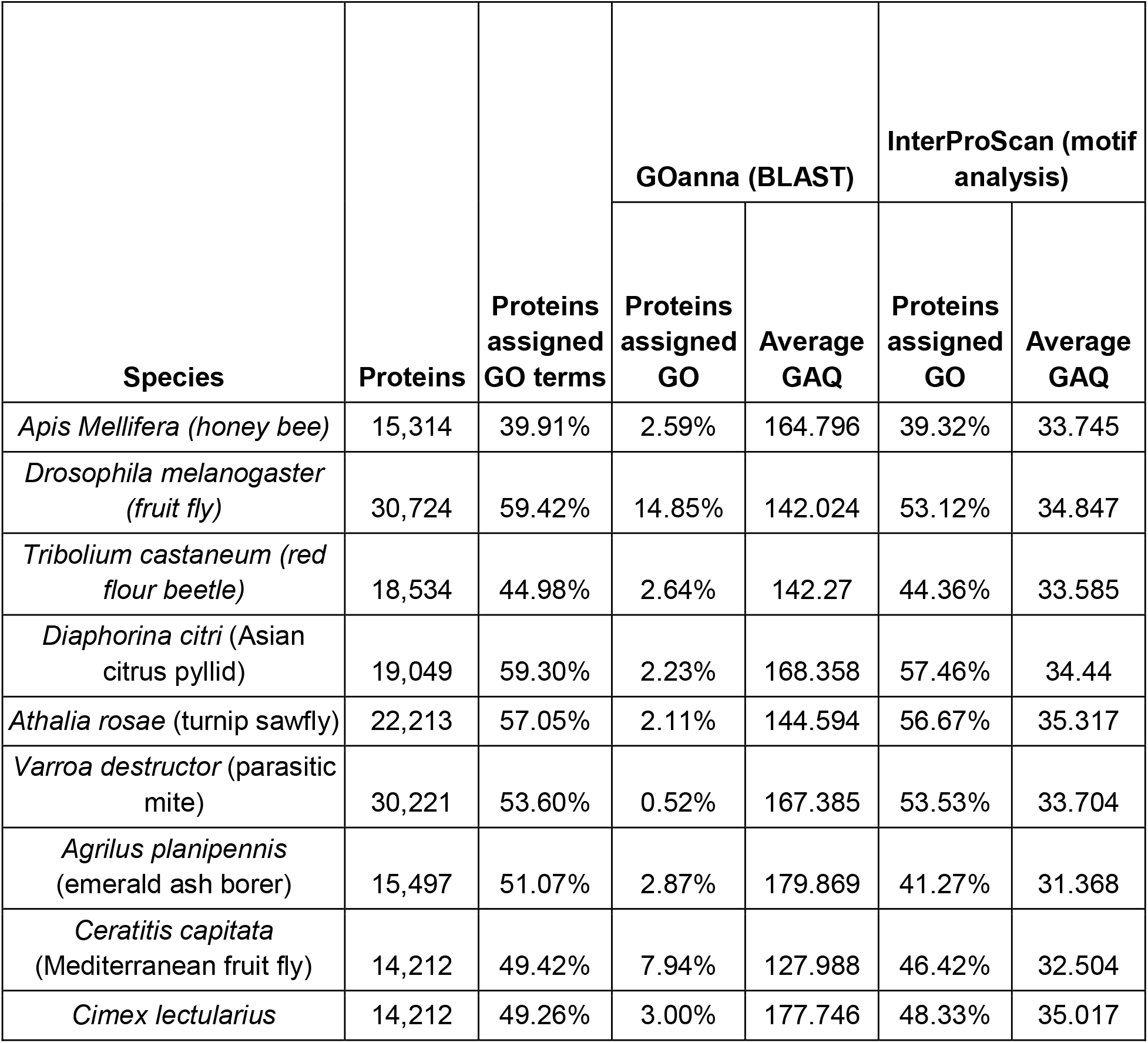
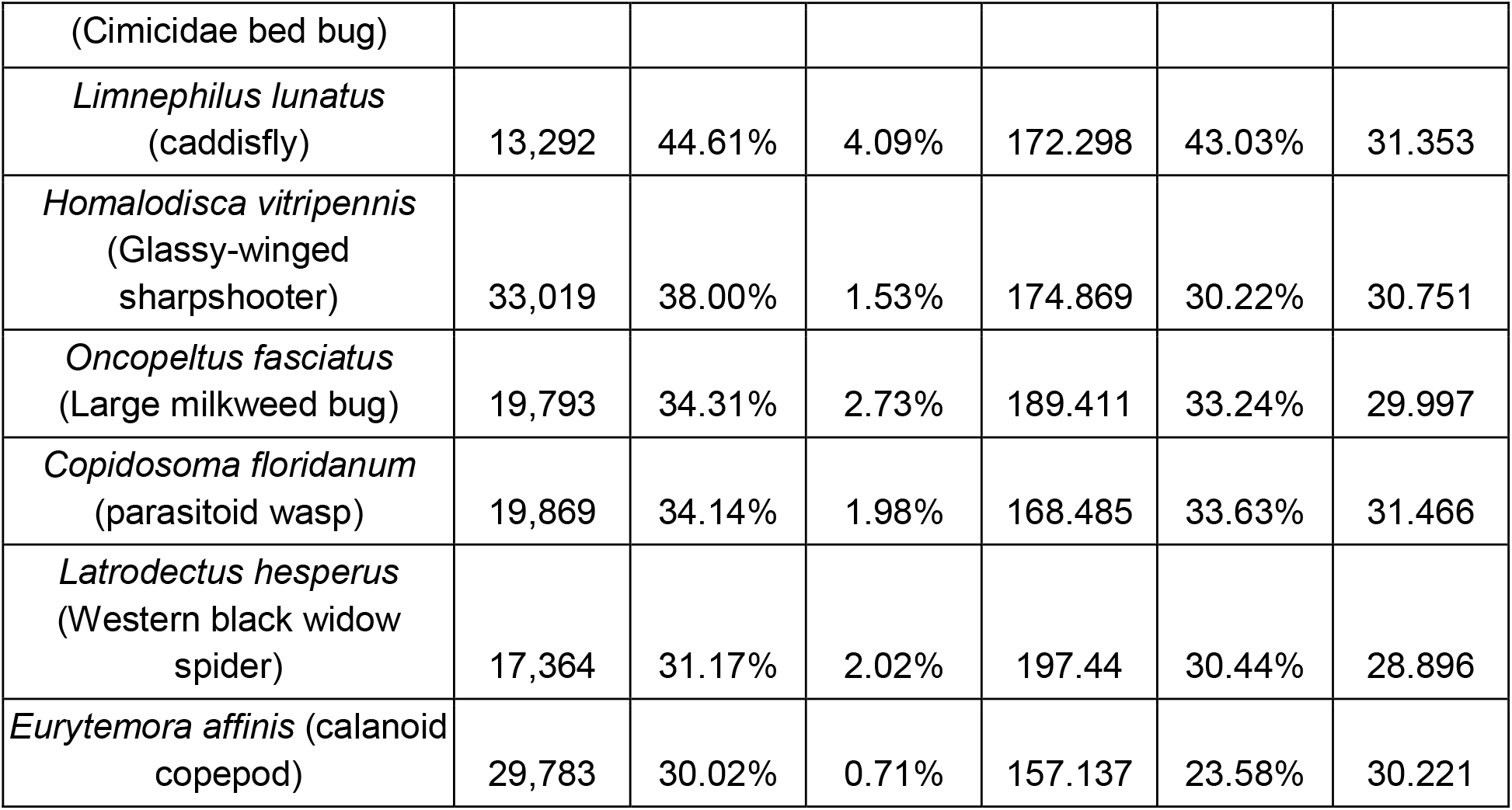
GOanna and InterProScan results for arthropod genomes selected for this study. The test species are sorted by their GO term coverage.

In addition to measuring how the assembly quality and proteome completeness influenced the GO term annotation, another question of interest was the potential influence of the phylogenetic distance from the model species, specifically *Drosophila melanogaster*. Among the reference genomes, *D. melanogaster* is by far the best annotated and curated. Since GOanna uses a database of experimentally validated GO terms wherein Drosophila was the model system used, 14.8% of *D. melanogaster* proteins were annotated with GO terms by GOanna compared to 2.5% and 2.6% for the honey bee and red flour beetle, respectively (Table 5).

Both *D. citri* (Asian citrus psyllid) and *V. destructor* (parasitic mite) showed overall annotation comparable to the selected references making the case that good quality genomes and annotation provide the best foundation for successful functional annotation. Surprisingly, the hymenopteran *A. rosae* (turnip sawfly) with a 99.7 BUSCO completeness, but lower contig N50 (51.4 kb) and scaffold N50 (1.3 Mb) than *D. citri* and *V. destructor* also fared well for overall annotation. The contiguous *D. citri* and *V. destructor* genomes did not have the highest BUSCO completeness scores (87.1% and 95.9%). The BUSCO ortholog set is computed based on a set of conserved genes in a clade and the hemipteran clade is relatively under-sampled among arthropods so this score might change in the future as more hemipteran genomes are sequenced.

*C. capitata* (Mediterranean fruit fly) had the highest percentage of proteins annotated by GOanna (7.9%), but that is somewhat expected considering its phylogenetic closeness to the reference species, *D. melanogaster*. The *L. lunatus* (caddisfly) and *L. hesperus* (Western black widow spider) genomes have the lowest contig N50, scaffold N50 metrics and BUSCO completeness scores but 44.6% of *L. lunatus* proteins were annotated compared to 31.17% of *L. hesperus* proteins. *E. affinis* (calanoid copepod) scored the poorest on GO annotation among out test species with only 30% of proteins annotated, possibly due to its phylogenetic distance from Drosophila, despite having a better contig and scaffold N50 of 5.7 kb and 862.6 kb respectively. However, it had a poor BUSCO completeness metric with only 57.5% completeness and 22.5% missing orthologs. We found a common theme in our test set and related analysis whereby the quality and depth of functional annotation was inversely proportional to the phylogenetic distance from the Drosophila model species (data not shown) . This emphasizes the need for better annotation of non-model species in every major clade so that proteins from newly sequenced genomes can be assigned function more accurately.

#### Pathway Annotation

High throughput sequencing has enabled the profiling of longitudinal transcriptional response at the organismal, tissue and single cell level in addition to multiple life stages and conditions. Although GO terms are highly effective at deducing the changes in gene expression, pathways-level perturbations provide valuable biological insight for the interpretation of functional genomics data sets and are critical for integrating proteome and metabolome data sets to understand phenotypes. Therefore, we were also interested in the ability to automatically reconstruct metabolic pathways from the proteomes from a range of arthropod genomes.

Pathways data is provided by resources such as KEGG [26], Reactome [27] and BioCyc [28] and as we developed our workflow, we selected KEGG pathways for our workflow because it supports the most extensive set of invertebrate species, and the KOBAS tool is freely available [23]. In our initial tests using the KOBAS tool to annotate pathways, we determined that comparing the arthropod proteome sets against the KEGG *Drosophila melanogaster* (‘dme’) provide the most comprehensive results, and this well-studied arthropod species also has the broadest set of functional information based on experimental validation, including pathways.

Not surprisingly, *A. mellifera* and *T. castaneum* references had similar proportions of proteins assigned to pathways although a slightly lower number of proteins per pathway than Drosophila (Table 6). The reference species had about one third of proteins assigned to pathways and most of the test species were annotated to the same degree or better. Curiously, several species did substantially better than the reference set: *V. destructor, A. rosae, D. citri* and *L. lunatus* all had about 40% of proteins assigned to pathways, and a similar effect was seen for the GO annotation in these species. We note that most of these species have well assembled genomes with a high contig and scaffold N50 and BUSCO completeness scores. The average number of proteins per pathway scaled with the genome contiguity and BUSCO duplication rate, suggesting that the higher gene copy number accounts for this variance (Supplementary Figure 1 and 2).

**Table 6.** KOBAS results for arthropod genomes selected for this study. The test species are sorted by the overall proportion of proteins assigned to pathways.

| Species | Proteins | All Pathways |  | KEGG Pathways |  |
| --- | --- | --- | --- | --- | --- |
|  |  | Proteins assigned to pathways | Average number of proteins in pathways | % assigned to pathways | Average number of proteins in pathways |
| *Apis mellifera* (Honeybee) | 15,314 | 29.27% | 3.41 | 17.57% | 20.23 |
| *Drosophila melanogaster* (fruit fly) | 30,724 | 37.73% | 8.77 | 21.24% | 49.08 |
| *Tribolium castaneum* (red flour beetle) | 18,534 | 30.03% | 4.22 | 16.99% | 23.68 |
| *Varroa destructor* (parasitic mite) | 30,221 | 41.55% | 9.63 | 23.50% | 54.62 |
| *Athalia rosae* (turnip sawfly) | 22,213 | 40.95% | 6.9 | 22.79% | 38.06 |
| *Diaphorina citri* (Asian citrus psyllid) | 19,049 | 40.07% | 5.88 | 23.72% | 34.75 |
| *Limnephilus lunatus* (caddisfly) | 13,292 | 38.09% | 3.92 | 22.94% | 23.10 |
| *Cimex lectularius* (Cimicidae bed bug) | 14,212 | 37.07% | 4.01 | 22.50% | 24.22 |
| *Ceratitis capitata* (Mediterranean fruit fly) | 12,318 | 35.91% | 3.35 | 21.36% | 19.78 |
| *Oncopeltus fasciatus* (Large milkweed bug) | 19,793 | 32.51% | 4.9 | 18.36% | 27.53 |
| *Agrilus planipennis* (emerald ash borer) | 15,497 | 31.81% | 3.74 | 18.92% | 22.05 |
| *Latrodectus hesperus* (Western black widow spider) | 17,364 | 30.06% | 4.06 | 16.97% | 22.66 |
| *Homalodisca vitripennis* (Glassy-winged sharpshooter) | 33,019 | 25.41% | 6.39 | 15.06% | 37.68 |
| *Copidosoma floridanum* (parasitoid wasp) | 19,869 | 25.35% | 3.83 | 14.43% | 21.56 |
| *Eurytemora affinis* (calanoid copepod) | 29,783 | 20.55% | 4.69 | 11.42% | 25.58 |

## Conclusion

Our results with a test set of arthropod genomes that are phylogenetically divergent and at different levels of assembly and annotation quality demonstrate the overall utility of our workflow to rapidly provide functional annotation for proteins. We are currently working on expanding functional annotation to include noncoding RNAs. Our workflow assigns GO and pathways information to 40-60% of proteins. While starting with a contiguous chromosomal length genome assembly and an evidence based protein set is ideal, we expect that species with complete gene models are sufficient to get a first-pass functional annotation. This functional information can be of immediate use to the community to support functional and comparative studies, including those generated by the Ag100Pest Initiative and other genomes hosted by the i5k Workspace@NAL. However, we would like to caution the user that the data sets underlying any functional annotation workflows are continually changing, and any functional annotation set should be refreshed periodically irrespective of whether or not the genome sequencing and annotation has changed. Furthermore, functional annotation provides information about pathways and gene families that are poorly annotated or absent from gene sets, providing useful information that can be used to direct targeted manual curation of genes. Manual curation of gene models is a well-established activity in the arthropod research community using Apollo [29] through community databases such as the i5k workspace@NAL [19], VectorBase [30], the Hymenoptera Genome Database [31], Citrus Greening Database [32–38], and others. Functional annotation would support this focus while extending the utility of the genome for the research community.

## Funding

This work was supported by the U.S. Department of Agriculture, Agricultural Research Service (USDA-ARS) and used resources provided by the SCINet project of the USDA-ARS, ARS project number 0500-00093-001-00-D. Mention of trade names or commercial products in this publication is solely for the purpose of providing specific information and does not imply recommendation or endorsement by the U.S. Department of Agriculture. USDA is an equal opportunity provider and employer.

## Acknowledgements

We would like to thank Lukas A. Mueller at Boyce Thompson Institute for providing computing facilities for data analysis.

## Author contributions

Conceptualization, F.M., A.K.C., M.F.P.; Methodology, F.M., A.M.C., S.S.; Software, A.M.C., S.S.; Formal Analysis, S.S.; Writing – Original Draft Preparation, F.M., M.F.P., S.S.; Writing – Review & Editing, A.K.C., A.M.C.; Visualization, S.S.; Project Administration, F.M., M.F.P.; Funding Acquisition, F.M., M.F.P.

## Data Availability Statement

The outputs from the workflow for each genome will be made available on AgData Commons. The docker containers are available at docker hub: GOanna, InterProScan, Combine GAFs and KOBAS (https://hub.docker.com/u/agbase). The source code for constructing the GOanna, InterProScan, Combine GAF and KOBAS containers is available on GitHub (https://github.com/AgBase/).

## Conflicts of Interest

The authors declare no conflict of interest.

## Supplementary data

**Supplementary Table 1.** GOanna version 2.2 parameters. Parameters are mainly based upon standard BLAST parameters and are categorized into required and optional. The parameters recommended for optimization are denoted with an *.

| Option | Description |
| --- | --- |
| **Required parameters** |  |
| -a* | BLAST database basename ('arthropod', 'bacteria', 'bird', 'crustacean', 'fish', 'fungi', 'human', 'insecta', 'invertebrates', 'mammals', 'nematode', 'plants', 'rodents', 'uniprot_sprot', 'uniprot_trembl', 'vertebrates' or 'viruses') |
| -c | Peptide FASTA filename |
| -o | BLAST output file basename |
| **Optional parameters** |  |
| -b* | Transfer GO with experimental evidence only ('yes' or 'no'). Default = 'yes'. |
| -d | Database of query ID. If your entry contains spaces either substitute and underscore (_) or, to preserve the space, use quotes around your entry. Default: 'user_input_db' |
| -e* | Expect value (E) for saving hits. Default is 10. |
| -f | Number of aligned sequences to keep. Default: 3 |
| -g | BLAST percent identity above which match should be kept. Default: keep all matches. |
| -h | Help |
| -m* | BLAST percent positive identity above which match should be kept. Default: keep all matches. |
| -s | Bit score above which match should be kept. Default: keep all matches. |
| -k* | Maximum number of gap openings allowed for match to be kept.Default: 100 |
| -l | Maximum number of total gaps allowed for match to be kept. Default: 1000 |
| -q* | Minimum query coverage per subject for match to be kept. Default: keep all matches |
| -r* | Ratio of query length to subject length. Lengths should be comparable for matches to be kept. Default: less than 1.2 so difference of up to 20% can be tolerated |
| -t | Number of threads. Default: 8 |
| -u | 'Assigned by' field of your GAF output file. If your entry contains spaces (eg. firstname lastname) either substitute and underscore (_) or, to preserve the space, use quotes around your entry (eg. firstname lastname) Default: 'user' |
| -x | Taxon ID of the peptides you are BLASTing. Default: 'taxon:0000' |
| -p | parse_deflines. Parse query and subject bar delimited sequence identifiers |

**Supplementary Table 2.** InterProScan version 5.45-80 parameters. The parameters are categorized into required and optional. The parameters recommended for optimization are denoted with an *

| Option | Description |
| --- | --- |
| Required parameters |  |
| -i | path to FASTA file that should be loaded on Master startup. Alternatively, in CONVERT mode, the InterProScan 5 XML file to convert. |
| Optional parameters |  |
| -a* | Comma separated list of analyses. If this option is not set, ALL analyses will be run. |
|  | Available analyses:<br>• TIGRFAM<br>• SFLD<br>• ProDom<br>• Hamap<br>• SMART<br>• CDD<br>• ProSiteProfiles<br>• ProSitePatterns<br>• SUPERFAMILY<br>• PRINTS<br>• PANTHER<br>• Gene3D<br>• Pfam<br>• Coils<br>• MobiDBLite |
| -b | Base output filename (relative or absolute path).<br>Note that this option, the output directory (-d) option and the output file name (-o) option are mutually exclusive. The appropriate file extension for the output format(s) will be appended automatically. By default the input file path/name will be used. |
| -d | Output directory.<br>Note that this option, the output file name (-o) option and the output file base (-b) option are mutually exclusive. The output filename(s) are the same as the input filename, with the appropriate file extension(s) for the output format(s) appended automatically. |
| -c* | Disables use of the precalculated match lookup service from EBI. All match calculations will be run locally. |
| -C | Supply the number of cpus to use. |
| -e | Excludes sites from the XML, JSON output |
| -f | Case-insensitive, comma separated list of output formats. Supported formats are TSV, XML, JSON, GFF3, HTML and SVG. Default for protein sequences are TSV, XML and GFF3, or for nucleotide sequences GFF3 and XML. |
| -g* | Switch on lookup of corresponding Gene Ontology annotation (IMPLIES -l lookup option) |
| -h | Display help information |
| -l | Also include lookup of corresponding InterPro annotation in the TSV and GFF3 output formats. |
| -m | Minimum nucleotide size of ORF to report. Will only be considered if n is specified as a sequence type. Please be aware of the fact that if you specify a too short value it might be that the analysis takes a very long time! |
| -o | Explicit output file name (relative or absolute path).<br>Note that this option, the output directory -d option and the output file basename -b option are mutually exclusive. If this option is given, you MUST specify a single output format using the -f option. The output file name will not be modified. Note that specifying an output file name using this option OVERWRITES ANY EXISTING FILE. |
| -p* | Switch on lookup of corresponding Pathway annotation (IMPLIES -l lookup option) |
| -t | The type of the input sequences (dna/rna (n) or protein (p)). The default sequence type is protein. |
| -T | Specify temporary file directory (relative or absolute path). The default location is temp/. |
| -v | Display version number |
| -r | 'Mode' required ( -r 'cluster') to run in cluster mode. These options are |
|  | provided but have not been tested with this wrapper script. For more information on running InterProScan in cluster mode:<br>https://github.com/ebi-pf-team/interproscan/wiki/ClusterMode |
| -R | Cluster run id (crid) required when using cluster mode. |
| -F | This is the output directory from InterProScan.(XML parser option) |
| -D | Supply the database responsible for these annotations. (XML parser option) |
| -x | NCBI taxon ID of the ID being annotated (XML parser option) |
| -y | Transcript or protein (XML parser option) |
| -n | Name of the biocurator who made these annotations (XML parser option) |
| -M | Mapping file (XML parser option) |
| -B | Bad input sequence file (XML parser option) |

**Supplementary Table 3.** KOBAS version 3.0.3 parameters. The parameters are categorized into required and optional. The parameters recommended for optimization are denoted with an *

| Option | Description |
| --- | --- |
| Required parameters |  |
| -i | INFILE can be FASTA or one-per-line identifiers. See -t intype for details. |
| -s* | SPECIES 3 or 4 letter species abbreviation (can be found here: ftp://ftp.cbi.pku.edu.cn/pub/KOBAS_3.0_DOWNLOAD/species_abbr.txt or here: https://www.kegg.jp/kegg/catalog/org_list.html) |
| -o | OUTPUT file (Default is stdout.) |
| -t | INTYPE (fasta:pro, fasta:nuc, blastout:xml, blastout:tab, id:ncbigi, id:uniprot, id:ensembl, id:ncbigene), default fasta:pro |
| -a or -g | -a runs KOBAS Annotate and -g runs KOBAS Identify. One of these options has to be used. Otherwise -j can be used to run both |
| Optional parameters |  |
| -l | LIST available species, or list available databases for a specific species |
| -e* | EVALUE expect threshold for BLAST, default 1e-5 |
| -r* | RANK rank cutoff for valid hits from BLAST result, default is 5 |
| -C* | COVERAGE subject coverage cutoff for BLAST, default 0 |
| -z* | ORTHOLOG whether only use orthologs for cross-species annotation or not, default NO (if only using orthologs, please provide the species abbreviation of your input) |
| -k | KOBAS HOME The path to kobas_home, which is the parent directory of sqlite3/ and seq_pep/. This is the absolute path in the container. |
| -v | BLAST HOME The path to blast_home, which is the parent directory of blastx and blastp. This is the absolute path in the container. |
| -y | BLASTDB The path to seq_pep/. This is the absolute path in the container. |
| -q | KOBASDB The path to sqlite3/, This is the absolute path in the container. |
| -p | BLASTP The path to blastp. This is the absolute path in the container. |
| -x | BLASTX The path to blastx. This is the absolute path in the container. |
| -T | number of THREADS to use in BLAST search. Default = 8 |
| -f | FGFILE foreground file, the output of annotate (KOBAS identify option) |
| -b | BGFILE background file, species abbreviation, see this list for species codes: https://www.kegg.jp/kegg/catalog/org_list.html (KOBAS identify option) |
| -d | DB databases for selection, 1-letter abbreviation separated by /: K for KEGG PATHWAY, n for PID, b for BioCarta, R for Reactome, B for BioCyc, p for PANTHER, o for OMIM, k for KEGG DISEASE, f for FunDO, g for GAD, N for NHGRI GWAS Catalog and G for Gene Ontology, default K/n/b/R/B/p/o/k/f/g/N/ (KOBAS identify option) |
| -m | METHOD choose statistical test method: b for binomial test, c for chi-square test, h for hypergeometric test / Fisher's exact test, and x for frequency list, default hypergeometric test / Fisher's exact test (KOBAS identify option) |
| -n | FDR choose false discovery rate (FDR) correction method: BH for Benjamini and Hochberg, BY for Benjamini and Yekutieli, QVALUE, and None, default BH (KOBAS identify option) |
| -c | CUTOFF terms with less than cutoff number of genes are not used for statistical tests, default 5 (KOBAS identify option) |

**Supplementary Figure 1:**
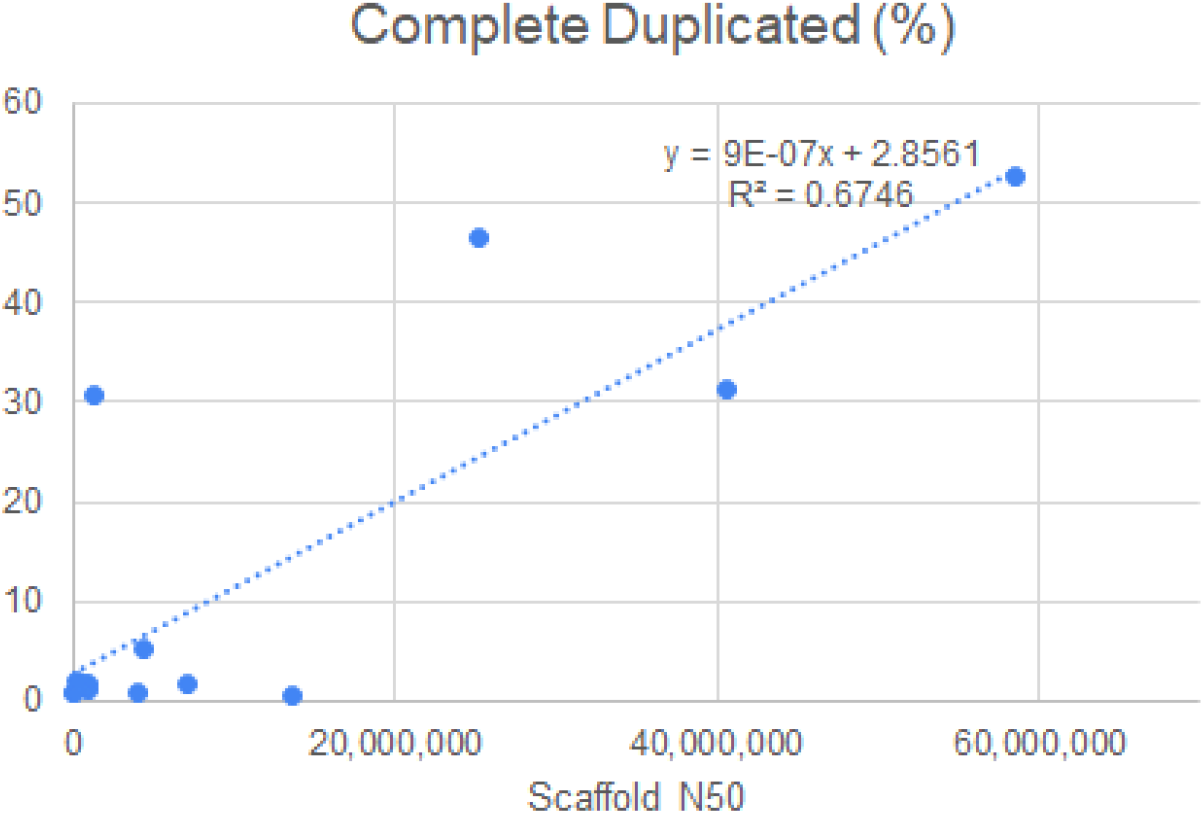
Duplication in BUSCO single copy orthologs: Plot of duplication (%) of 1013 single copy orthologs against the scaffold N50 showing correlation of increasing duplication with increase in contiguity of the assembly.

**Supplementary Figure 2:**
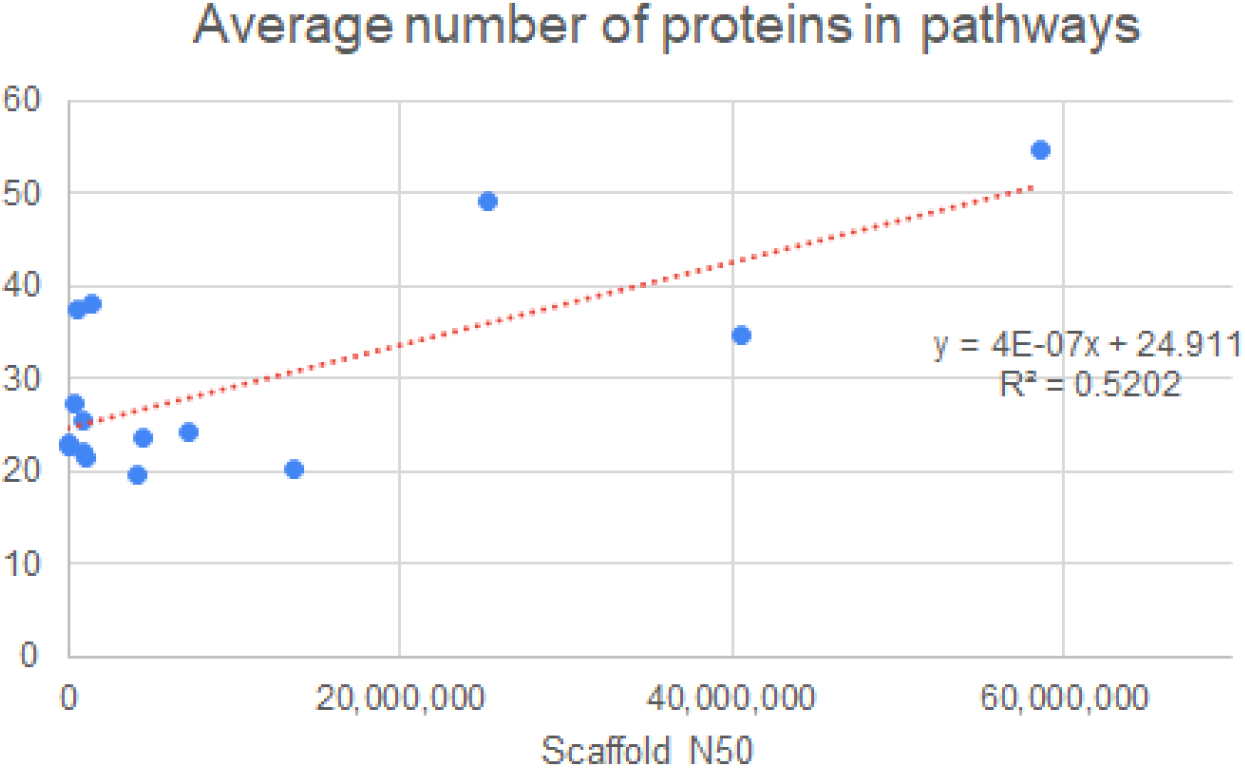
Average number of proteins per pathway: Plot of average number of proteins per pathway against the scaffold N50 showing a correlation of increasing protein count with increase in contiguity of the assembly.

## Notes

https://agbase-docs.readthedocs.io/en/latest/agbase/workflow.html

### Competing Interest Statement

The authors have declared no competing interest.

